# DARTS: an Algorithm for Domain-Associated RetroTransposon Search in Genome Assemblies

**DOI:** 10.1101/2021.12.03.471067

**Authors:** Mikhail Biryukov, Kirill Ustyantsev

## Abstract

Retrotransposons comprise a substantial fraction of eukaryotic genomes reaching the highest proportions in plants. Therefore, identification and annotation of retrotransposons is an important task in studying regulation and evolution of plant genomes. A majority of computational tools for mining transposable elements (TEs) are designed for subsequent genome repeat masking, often leaving aside the element lineage classification and its protein domain composition. Additionally, studies focused on diversity and evolution of a particular group of retrotransposons often require substantial customization efforts from researchers to adapt existing software to their needs. Here, we developed a computational pipeline to mine sequences of protein-coding retrotransposons based on the sequences of their conserved protein domains - DARTS. Using the most abundant group of TEs in plants - long terminal repeat (LTR) retrotransposons (LTR-RTs), we show that DARTS has radically higher sensitivity of LTR-RTs identification compared to a widely accepted LTRharvest tool. DARTS can be easily customized for specific user needs. As a result, DARTS returns a set of structurally annotated nucleotide and amino acid sequences which can be readily used in subsequent comparative and phylogenetic analyses. DARTS should facilitate researchers interested in discovery and in-detail analysis of diversity and evolution of retrotransposons, LTR-RTs, and other protein-coding TEs.

## Introduction

Transposable elements (TEs) are important players in the evolution of genomes [1–4]. TEs activity drives genetic diversity, contributes to establishment of new gene regulatory networks and rewiring of the existing ones, as well as can result in the origin of new genes sequestered by the host genome for its functioning [5–7]. Long-term existence and evolution of TEs resulted in a broad diversity of the mechanisms for their transposition and replication, and origin of a variety of different structural variants [8,9].

Retrotransposons, a group of TEs which move through a reverse transcription mechanism, are the most ubiquitous TEs in eukaryotic genomes. Due to their propensity to increase in copy number, retrotransposons constitute a substantial portion of the host genomes reaching as high as 80% of the total genome size in some plants [10,11]. Thus, studying retrotransposons is an essential part of understanding plant evolution. The majority of retrotransposons in plants are long terminal repeat (LTR) retrotransposons (LTR-RTs), which are structurally and evolutionarily similar to retroviruses of vertebrates [12,13]. Autonomous, i. e. capable of self replication, LTR-RTs are complex genetic entities consisting of several protein-coding domains and non-coding regulatory sequences (such as LTRs), which mediate transcription, replication, and integration of the TEs [14–17]. Despite similarities in the general replication mechanism, LTR-RTs are structurally diverse and encode for additional protein domains, which are supposed to fine-tune their life cycle [18–20]. Despite the structural differences, the central functional domain of all autonomous retrotransposons, reverse transcriptase (RT), remains conserved through evolution, allowing unbiased phylogenetic delineation and classification of retrotransposon diversity [9]. Evolution of LTR-RTs and other retrotransposons as individual entities attracts attention by itself being an example of modular evolution [21,22]. Modular evolution means that the main driving force is not a random mutational process, but acquisition, reshuffling, and loss of whole structural elements, such as protein domains and transcriptional enhancers. Therefore, the history of a distinct protein domain in a retrotransposon can be different from the evolution of its core RT domain [21,23,24].

The majority of computational tools developed for annotation of LTR-RTs in the genomic sequences initiate their search from identification of LTRs, and not conserved protein domains [25–27]. Alternative approaches, like RepeatModeler, first look for any repetitive sequence on the nucleotide sequence level and then try to classify them based on the homology information to known TEs [28]. However, in cases when it is important to search for a specific family of TEs, these methods, apart from being too redundant and computationally time-consuming, may end up in a very high rate of false-negative results, since some TEs lineages may be present in a very low number of copies, and some LTR-RTs copies may lack well-detectable LTRs. On the other hand, homology-based approaches suffer from incompleteness of the reference databases [29].

Here, based on our experience in retrotransposon identification [21,23], we developed a new computational tool which takes advantage of the conserved nature of protein domains encoded by retrotransposons - DARTS (an Algorithm for Domain-Associated RetroTransposon Search in Genome Assemblies). DARTS uses a constantly updating database of conserved protein domain sequences, which allows to mitigate issues of the incomplete reference databases and to customize it for identification of virtually any protein-coding group of TEs. Importantly, DARTS simultaneously performs structural annotation and extraction of sequences of the corresponding protein domains, which makes it a handy tool for studies of phylogeny and diversity of retrotransposons and other protein-coding TEs.

## Methods

### Data Collection

For the analysis, we downloaded genome reference assemblies from the NCBI Genome database (https://www.ncbi.nlm.nih.gov/genome/, accessed on 23 November 2021) of four model plant species: *Arabidopsis thaliana* (TAIR10.1, 120 Mbp), *Nicotiana tabacum* (Ntab-TN90, 3736 Mbp), *Selaginella moellendorffii* (GCF_000143415.4, 212 Mbp), and *Zea mays* (Zm-B73-REFERENCE-NAM-5.0, 2192 Mbp).

### Description of the DARTS pipeline

The DARTS pipeline consists of several scripts written in Python (v 3.6) and Bash programming languages. The scripts, installation and detailed usage manuals are available on GitHub: https://github.com/Mikkey-the-turtle/DARTS_v0.1. The general pipeline scheme is shown in Figure 1. A user may choose which parts of the pipeline to execute and can customize every filtering and threshold values presented in the default version which was originally adapted for identification of LTR-RTs.

**Figure 1.**
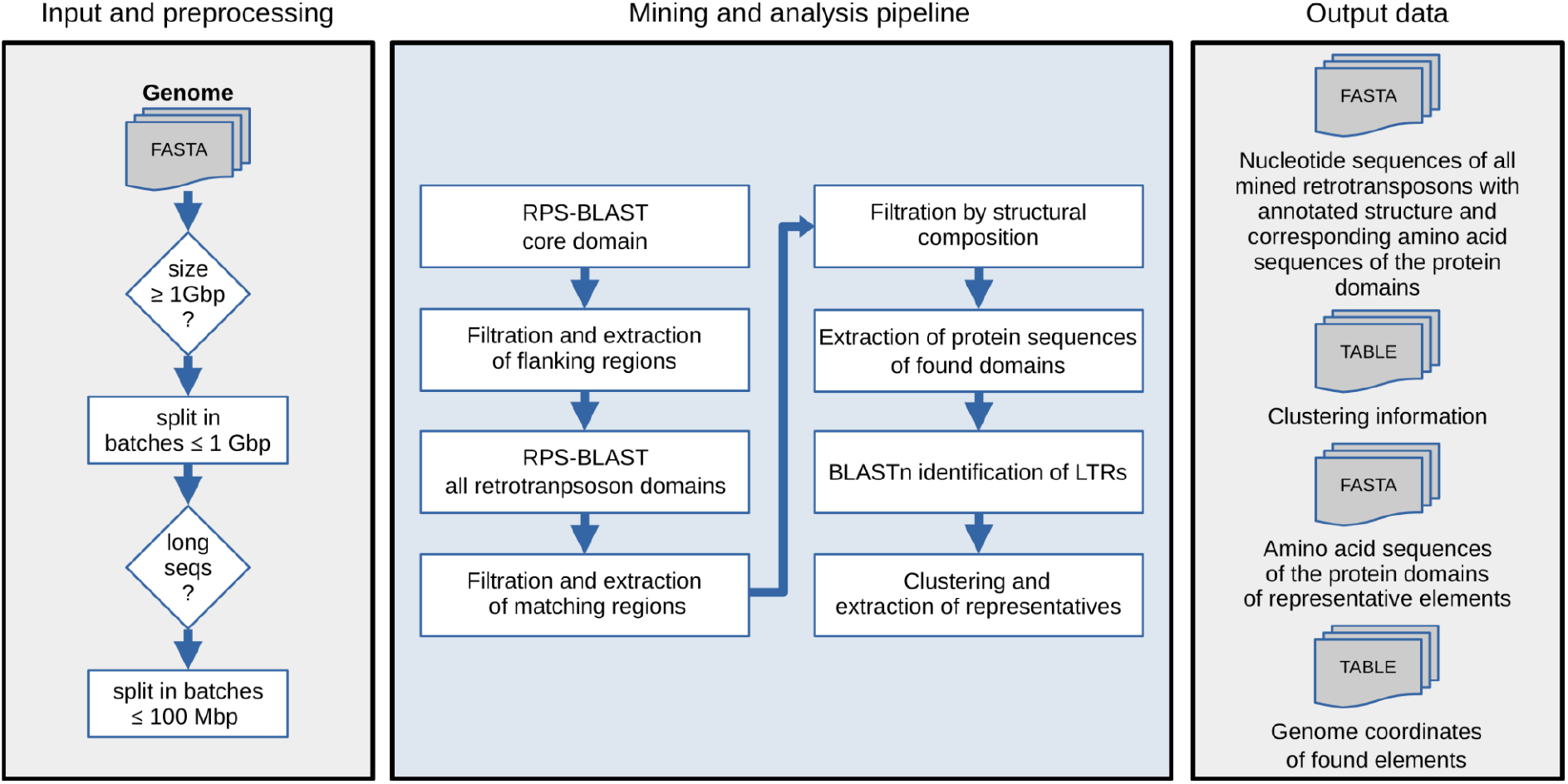
Principle scheme of the DARTS workflow. Detailed description of each step is in the text.

Before the analysis, the program checks the genome size, and splits it into several batches below 1 Gbp to allow fast search. If the genome assembly consisted of long sequences such as fully-assembled chromosomes, the sequences were split into batches below 100Mbp. To identify target protein domains, DARTS uses standalone Reverse PSI-BLAST (RPS-BLAST) from the BLAST+ package [30] supplemented with corresponding multiple sequence alignment protein models, or profiles, obtained from a local copy of NCBI Conserved Domain Database (CDD) [31] (https://www.ncbi.nlm.nih.gov/Structure/cdd/cdd.shtml, Accessed on November 2019). In the first RPS-BLAST search round, a single CDD profile of interest (e.g. reverse transcriptase domain specific to LTR-RTs) is used to scan the genomic sequences. This results in a set of matches which are pre-filtered by *e-value* (1e-3) and length of the match. Genomic coordinates of a match are identified, and the corresponding nucleotide sequence with flanking regions of length 7500 bp is extracted for each match. The second RPS-BLAST search utilizes a database of all CDD profiles, including the first profile, which are expected to be found in the extracted sequence regions. Processing of the second RPS-BLAST run results in structural annotation of TEs of interest and subsequent filtration of the elements by presence of a user-defined set of protein domains. Importantly, when several core domains are present in the same expanded matching region, DARTS will try to delineate them into separate domain assemblies. When a domain match is interrupted by frameshifts or small insertions, DARTS will assemble its parts in a single unit for annotation (Figure S1). Amino acid sequences of each of the identified domains are extracted and stored in separate FASTA-formatted files. For LTR-RTs, using the BLASTn tool from the BLAST+ package [30], DARTS will attempt to identify LTRs flanking the first and the last identified protein domains with more than 80% identity and more than 100 bp but less than 3000 bp in length. Each element obtains a score (%score) based on the number of identified protein domains, length and quality of the matches, presence of uninterrupted open reading frames (ORFs), and LTRs identity for LTR-RTs. Each sequence which passed the filtration will have a unique name identifier presented in the following format: “%project_name_%batch_%num_ID|%structure|%LTR_information|%score”, where %project_name - user-defined name of the DARTS run, %batch - number of the corresponding genome batch-file, %num_ID - numerical identifier in the current genome batch-file, %structure - generalized protein domain-based structure presented for *Ty3/gypsy* LTR-RTs (e.g. “GAG.PRo.gRT.gRH.INT”), %LTR_information is shown as LTR%identity-length (e.g. LTR%99.567-232), %score - float number.

For the purpose of subsequent comparative and phylogenetic analyses, DARTS can reduce redundancy of the dataset through clustering using MMseqs2 [32] and subsequent selection of clusters’ representatives based on the %score value and structural composition. Clustering information is stored as a tab-separated values table file and can be later reanalyzed using custom criteria. For LTR-RTs, by default clustering is performed using the core and the most conserved domain, reverse transcriptase (RT), with the following default parameters: “easy-cluster -min-seq-id 0.8 -c 0.8”. Nucleotide sequences of the representative elements, and amino acid sequences of each of their protein domains are deposited in separate FASTA-formatted files. These sequences can later be directly used for multiple sequence alignment generation for subsequent phylogenetic analysis.

### Identification of LTR retrotransposons using LTRharvest

To mine LTR-RTs from the selected plant genomes using de novo LTR-RTs prediction tool LTRharvest [26], we ran the program with the following parameters: “-minlenltr 200, -maxlenltr 2000, -mindistltr 3000, -maxdistltr 22000, -similar 85.0, -overlaps no, -mintsd 3, -maxtsd 20”. The resulting file with all hypothetical nucleotide full-length LTR-RTs sequences produced by LTRharvest was then processed by DARTS to identify sequences containing the RT domain and to ensure unbiased comparison between both tools. To compare numbers of elements uniquely identified by both the DARTS and LTRharvest tools, we performed reciprocal BLASTn searches with “-max_target_seqs 1” parameter.

## Results and Discussion

### Application of DARTS for general and lineage-specific LTR retrotransposon identification

Previously, we performed a study on diversity and evolution of a structurally variable group of *Ty3/gypsy* plant LTR-RTs - *Tat* [23,33,34]. *Tat* LTR-RTs have an additional ribonuclease H domain (aRNH) of the so-called archaeal origin which is fixed in several positions with regard to other domains in different *Tat* lineages [23]. In our previous study on *Tat* [23], we used a conventional tool for de novo prediction of LTR-RTs - LTRharvest [26]. Later, doing an independent search using tBLASTn with aRNH sequence as a query we found out that a substantial fraction of aRNH-containing *Tat* LTR-RTs was underrepresented in the LTRharvest output. We reasoned that this can be explained by the majority of LTR-RTs copies in the studied plant genomes being damaged, fragmented (not intact), and lacking detectable LTRs. The fact that LTRs are used as a starting point for LTR-RTs identification in several published software [25–27,35], including LTRharvest (now a part of the most popular de novo repeat identification pipeline RepeatModeler [28]), inspired us to develop a new algorithm that could automatically perform identification of protein-coding TEs and LTR-RTs in particular. We named it Domain-Associated RetroTransposon Search (DARTS) as initiation of the screen and subsequent structural annotation are based on prediction of conserved protein domains and not LTR sequences. A basis for DARTS is our experience in semi-automated identification of both protein-coding LTR-RTs and non-LTR retrotransposons [21], and conceptually similar approaches performed by other researchers [20,36,37].

For the analysis, we selected reference genome assemblies of four widely used model plant species varying in the genome size and TEs content (see *2.1 Data Collection*) and applied DARTS and LTRharvest to identify LTR-RTs. The DARTS search was initiated with the most conserved reverse transcriptase (RT) domain, while LTRharvest attempts to identify regions flanked by direct repeat sequences of hypothetical LTRs [26]. Using DARTS, we mined 267,105 LTR-RTs elements (88,389 with LTRs) in the four studied genomes, while only 34,030 sequences predicted by LTRharvest contained the RT domain sequence after filtration of originally predicted 55,658 elements (Figure 2A). Importantly, all the 34,030 LTRharvest elements were also predicted by DARTS (Table S1), suggesting almost eight times higher sensitivity of the latter.

**Figure 2.**
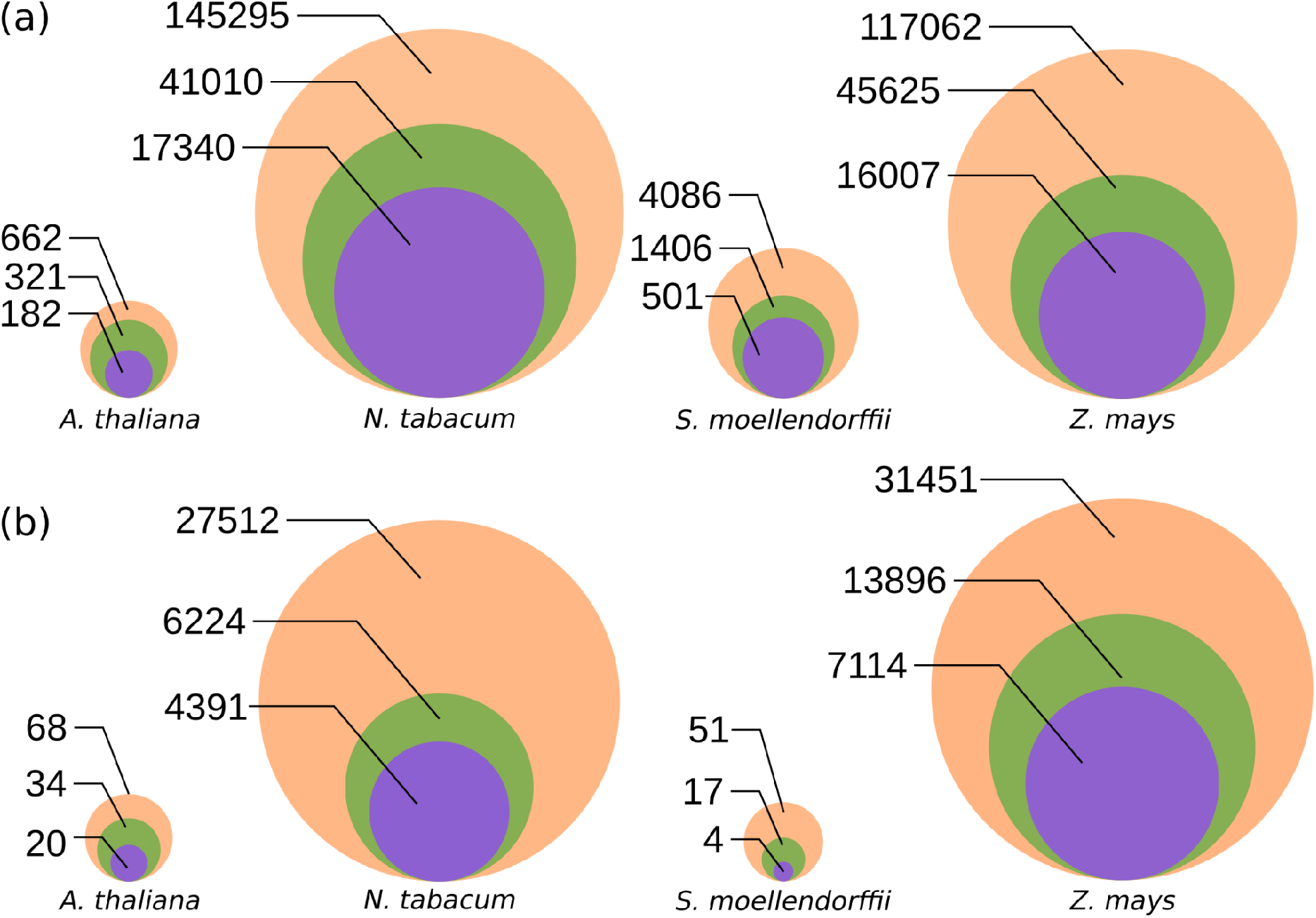
Sensitivity on LTR retrotransposon (LTR-RTs) identification by the DARTS and LTRharvest pipelines. Orange circles - number of elements found by DARTS; green circles - number of LTR-RTs by DARTS with predicted LTRs; purple circles - number of elements found by LTRharvest. Sizes of the circles are proportional to the number of elements with relation to the minimum and the maximum values. Exact numbers of elements are indicated to the left from the circles. Parameters of the LTRharvest search were the same for all the approaches (**a-b**). (**a**) Prediction of LTR-RTs by DARTS when the search is initiated from the RT domain; LTRharvest elements were retained if the RT domain was present. (**b**) Prediction of LTR-RTs by DARTS when the search is initiated from the aRNH domain; DARTS and LTRharvest elements were retained if both the RT and aRNH domains were present.

To exemplify DARTS performance when search is initiated from a different protein domain, we screened for *Tat* LTR-RTs in the same genome assemblies using the aRNH CDD profile in the first round of RPS-BLAST. DARTS found 59,082 elements (20,171 with LTRs) while only 11,529 LTR-RTs were identified by LTRharvest (Figure 2B). This suggests that the overall abundance of *Tat* LTR-RTs in our previous study where we used LTRharvest [23] was largely underestimated.

It must be noted that potential false-positive matches can be present when only the initial target domain is found. However, the chances of this are low since during the second step of the RPS-BLAST search all the domains are reannotated again which results in increase of *e-value* since the size of the database is decreased to a single sequence region. Nevertheless, the false-positive hits can be filtered out on the way to phylogenetic analysis standing as outliers during clustering and multiple sequence alignment compared to true-positive representatives. Alternatively, whenever it is possible, we would suggest to filter the results of DARTS by presence of one or two additional domains or regulatory sequences like LTRs to completely avoid the problem. In this study, we found that the number of RT-only containing matches in the RT domain search initiated by DARTS equals 5.8% ± 2.5% (mean ± standard deviation of the mean). Therefore, this range can be considered as a theoretical stringent upper boundary for the false-positive TEs detected by DARTS.

### Possible application of DARTS for identification of other transposable elements

Although here we show examples of DARTS usage for general LTR-RTs identification and targeted *Tat* LTR-RTs mining in plants, our primary object of interest, the software can be easily customized for search of other TEs with conserved protein-coding domains in other eukaryotic genomes. For example, Penelope-like retroelements can be targeted by search for their specific RT and endonuclease domains [38,39] and DIRS-like retrotransposons by the RT and tyrosine recombinase domains [40,41]. Various non-LTR retrotransposon groups apart from their specific RT domain have two types of endonucleases and two types of the RNH domains [23,37,42]. Cut-and-paste DNA transposons can be identified by the transposase domains, DNA helicases can be found in *Helitrons* and DNA polymerases in *Mavericks* [43–45].

For non-LTR retrotransposons and DNA transposons, DARTS initial identification approach is similar to the methods implemented in previously published software such as MGEScan-non-LTR and TransposonPSI [36,46]. However, DARTS is more advantageous since it can also perform automatic structural annotation and clustering, and its algorithm relies on constantly updating RPS-BLAST and the CDD database. Therefore, more sensitive profiles can be used to provide a detailed and targeted annotation of TEs of interest. Additionally, compared to TransposonPSI, DARTS returns amino acid sequences of each of the identified domains without need for additional parsing allowing direct transition to phylogenetic analysis.

A part of the TEs analysis that is not covered by DARTS is annotation of non-protein coding genes and copies lacking a domain of interest that was used to initiate the search. While annotation of such elements as SINEs and MITEs indeed requires a substantially different approach for their identification [28,47], severely damaged copies and sequences such as solo-LTRs lacking a domain of interest can still be found by applying nucleotide BLAST or RepeatMasker [30,48] using the copies identified by DARTS as queries. Thus, their number can be accounted for the genome annotation.

## Conclusions

Here, we developed a new pipeline for automatic search and structural annotation of protein-coding LTR-RTs and other retrotransposons in genomic sequences. DARTS is beneficial when one is interested in analysis of all the structural diversity of a TEs group. We showed that DARTS is almost eight times more sensitive in LTR-RTs identification than a de novo tool LTRharvest which is now included in a widely used RepeatModeler version 2 pipeline [28]. Easiness of the DARTS customization should facilitate many researchers studying diversity and evolution of different groups of TEs.

## Author Contributions

Conceptualization, M.B. and K.U.; formal analysis and software, M.B.; writing—original draft preparation, M.B. and K.U.; writing—review and editing, M.B. and K.U.; supervision, K.U.; funding acquisition, M.B. and K.U. All authors have read and agreed to the published version of the manuscript.

## Acknowledgements

We thank Prof. Dr. Eugene Berezikov for his valuable comments on the manuscript draft.

## Data Availability

The software and datasets produced in this study are openly available on GitHub https://github.com/Mikkey-the-turtle/DARTS_v0.1

## Captions for the supplementary files

**Table S1.** BLASTn matches between the LTR-RTs predicted by LTRharvest against the DARTS LTR-RTs matches.

**Figure S1**. A principle scheme of fragmented protein domain assembly by DARTS.

