## Supplementary material for "DARTS: an Algorithm for Domain-Associated RetroTransposon Search in Genome Assemblies": Figure S1

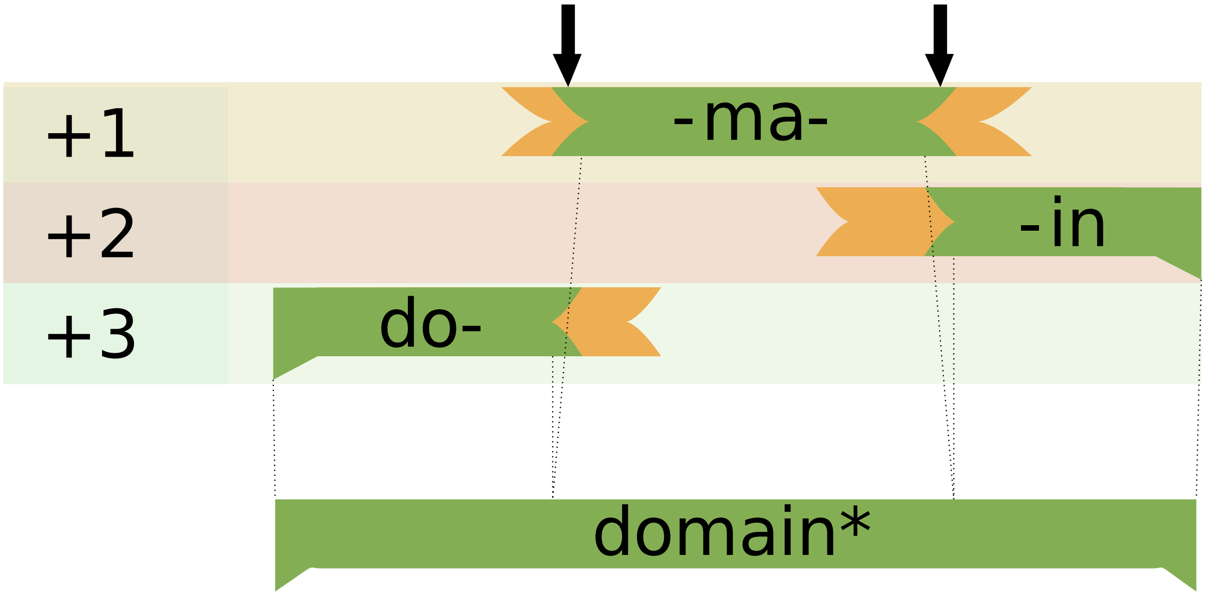


**Figure S1**. A principle scheme of fragmented protein domain assembly by DARTS. Translation frames are indicated as +1, +2 and +3. Black arrows indicate coordinates for merging of the fragmented domain sequences. Green regions are parts of the sequence with homology to the searched protein profile. Brown regions are non-homologous parts of the sequence. domain* - reconstructed sequence of the protein domain.
